# Impact of the inoculum size on the *in vivo* activity of the aztreonam-avibactam combination in a murine model of peritonitis due to *Escherichia coli* expressing CTX-M-15 and NDM-1

**DOI:** 10.1101/2024.09.18.613740

**Authors:** Laura Benchetrit, Ariane Amoura, Samuel Chosidow, Alice Le Menestrel, Victoire de Lastours, Françoise Chau, Sara Dion, Laurent Massias, Bruno Fantin, Agnès Lefort

**Affiliations:** IAME, UMR1137, Université Paris Cité, F-75006 Paris; Laboratoire de toxicologie, Hôpital Bichat, F-75018 Paris; Service de Médecine Interne, Hôpital Beaujon, F-92110 Clichy

## Abstract

**Background:** The combination of aztreonam (ATM) and avibactam (AVI) is an attractive option to treat infections caused by extended spectrum *β*-lactamase plus NDM-1-producing *Enterobacteriaceae*. Since ATM activity was shown to be severely impacted by an increase in the inoculum size *in vitro*, we wondered whether ATM-AVI activity could be impaired in high-inoculum infections.

**Methods:** We analyzed the impact of the inoculum size on ATM-AVI activity *in vitro* and in a murine model of peritonitis due to susceptible *E. coli* CFT073-pTOPO and its isogenic derivatives producing NDM-1 (*E. coli* CFT073-NDM1) and CTX-M-15 plus NDM-1 (*E. coli* CFT073-CTXM15-NDM1). The impact of the inoculum size on bacterial morphology was studied by microscopic examination.

**Results:** *In vitro*, at standard (10^5^) inoculum, *E. coli* CFT073-CTXM15-NDM1 was resistant to ATM but susceptible to the ATM-AVI combination. At high (10^7^) inoculum, MICs of ATM alone and of the ATM-AVI combination reached > 512 and 64 mg/L respectively, against all tested strains. ATM led to bacterial filamentation when active against the bacteria, i.e., in monotherapy or in combination with AVI against susceptible *E. coli* CFT073-pTOPO, and only in combination with AVI against *E. coli* CFT073-CTXM15-NDM1. *In vivo*, increase in the inoculum led to a drastic decrease in the activity of ATM alone against *E. coli* CFT073-pTOPO, and of ATM-AVI against *E. coli* CFT073-CTXM15-NDM1.

**Conclusion:** Our results suggest a high *in vivo* impact of the inoculum increase on the activity of ATM alone against ATM-susceptible *E. coli*, and of ATM-AVI against CTX-M-15 plus NDM-1 producing *E. coli*. Clinicians must be aware of the risk of failures when using AZT-AVI in high inoculum infections.

## Introduction

The massive consumption of carbapenems to treat infections caused by extended spectrum *β*-lactamase-producing *Enterobacteriaceae* (ESBL-PE) has contributed to the emergence of carbapenemase-producing strains (CPE) whose rapid spread represents a worrisome threat worldwide^1^. The major concern comes from metallo-β-lactamases (MBLs) which are endemic in many areas, such as India or Greece, because they confer resistance to nearly all available antibiotics, including novel *β*-lactams, leading to a therapeutic deadlock.

Aztreonam (ATM) is a natural *β*-lactam discovered in the 1980s. It is effective against gram-negative bacilli such as *Enterobacteriaceae* and *Pseudomonas aeruginosa*^2^. The pharmacokinetic/pharmacodynamics (PK/PD) index that best predicts its antimicrobial activity is the percentage of time in a 24-hour period when the free drug concentration is above the minimal inhibitory concentration (MIC) (*%f*T _> MIC_)^3^. An inventory has identified it as an old molecule worth studying thanks to its efficacy against MBLs^4^. Indeed, ATM is the only *β*-lactam effective against MBLs. However, its efficacy against MBL-producing isolates is most of the time compromised by the co-production of serine-protease enzymes of Ambler’s *β*-lactamases class A, C and D, that hydrolyze ATM^3,5^. In addition, ATM activity was shown to be greatly impacted by the size of the inoculum, so-called “inoculum effect” *in vitro*, although the *in vivo* impact of this effect on *Enterobacteriaceae* bearing MBL remains unknown,^6–8^.

Avibactam (AVI) is a semi synthetic non *β*-lactam *β*-lactamase inhibitor belonging to the diazabicyclooctanes^9^. It is effective against serine-protease enzymes of Ambler’s class A, C and D, by covalently acylating its serine target^10,11^. AVI alone does not display any intrinsic antimicrobial activity^12^. PK/PD parameter that best correlates with the efficacy of AVI in combination with ATM is the percentage of time in a 24-hour period when the free drug concentration is above the critical threshold of 2.5 mg/L (%*f*T _CT > 2,5 mg/L_)^9,13^.

The idea of combining ATM and AVI was hence put forward as AVI is stable against ESBL-PE while ATM is effective against MBLs such as NDM-1 enzyme^14^. The *in vitro* efficacy of ATM-AVI has been demonstrated^14^ but an inoculum effect of the combination was evidenced^15–18^. A few PK/PD studies on murine models supported the benefit of the combination^19,20^.

Some patients have already been successfully treated by ATM and ceftazidime-AVI (CZA) because ATM-AVI was not available for human use, ^21–24^. Recent phase 2a and phase 3 studies investigated pharmacodynamics and safety of ATM-AVI in patients with various infections^25,26^.

Thus, IDSA recommended using CZA and ATM to treat MBL-producing *Enterobacteriaceae* although extra data is necessary especially regarding the *in vivo* impact of the inoculum effect^27^.

Therefore, the aims of our study were to assess the efficacy of ATM-AVI combination against an NDM-1 plus CTX-M-15 producing *E. coli* strain, by comparison to its activity against *β*-lactam susceptible parental strain, in a murine model of peritonitis at standard and high inoculum.

## Material and methods

### Bacterial strains and plasmids

To conduct our experiments, 3 isogenic strains were constructed. Details of construction have already been reported^28^. Briefly, susceptible uropathogenic *E. coli* CFT-073 (O6:K2:H1)^29^ was used as recipient for the plasmid pCR-Blunt II-TOPO (Life Technologies, Saint-Aubin, France) carrying a kanamycin resistance gene. Regions corresponding to *bla*_NDM-1_ and *bla*_CTX-M-15_ were cloned in the pCR-Blunt II-TOPO resulting in plasmids pTOPO-NDM1 and pTOPO-CTXM15-NDM1. These plasmids were introduced by electrotransformation into *E. coli* CFT073, as described previously, resulting in *E. coli* CFT073-pTOPO, *E. coli* CFT073-NDM1 and *E. coli* CFT073-CTXM15-NDM1. Strains and subcultures were grown in Mueller Hinton (MH) broth using kanamycine (400 mg/L) to avoid all plasmid loses.

### Antimicrobial agents

Antibiotics used were kanamycin (Sigma-Aldrich, Saint-Quentin-Fallavier, France), ATM (Sigma-Aldrich, Saint-Quentin-Fallavier, France for *in vitro* experiments; Sanofi, Gentilly, France for *in vivo* experiments), AVI (Sigma-Aldrich, Saint-Quentin-Fallavier, France for *in vitro* experiments; MedChemEpress, Sollentuna, Sweden, for *in vivo* experiments) and imipenem (IPM) (Arrow, Lyon, France).

### *In vitro* experiments

#### Growth rate and fitness

Growth curves in Luria-Bertani medium were performed by measuring the OD at 600 nm every 5 minutes during 24 hours at 37°C using an automatic spectrophotometer (Tecan Infinite F200PRO, Männedorf, Switzerland). Results were analyzed using R software which provided maximum growth rates (MGR) used to compare bacterial fitness^30^.

#### MICs

MICs of ATM, AVI, ATM in the presence of a fixed concentration of 4 mg/L AVI (as recommended by the EUCAST for ATM plus AVI MIC determination), and IPM were determined by the micro dilution method in Mueller Hinton (MH) broth in accordance with EUCAST^31^.

To evaluate the impact of the inoculum on antibiotic activity, MICs were performed at standard (5×10^5^ CFU/mL) and high (5×10^7^ CFU/mL) inoculum. To analyze whether the results were dependent of the method used for MIC determination, MICs of ATM against *E. coli* CFT073-pTOPO were also determined by the agar dilution method of Steers *et al*., at standard (5×10^5^ CFU per spot) and high (5×10^7^ CFU per spot) inoculum^32^.

The impact of antibiotic exposure on bacterial morphology was analyzed before and after 24h antibiotic exposure using an optical microscope (Zeiss, Oberkochen, Germany) using a 100x objective after Gram staining.

All experiments were performed at least three times.

#### Time kill curves at standard and high inoculum and selection of mutants

For time kill curves, a culture of each strain was grown overnight in MH containing kanamycin (400 mg/L). The culture was diluted 1/1000 and incubated for 4h30 to obtain an exponential phase culture of 10^7^ to 10^8^ CFU/mL for the high inoculum assay. This culture was diluted 1/100 for the standard inoculum assay. ATM was added at 40 mg/L, four times the residual free drug concentration in infected mice, and AVI at 4 mg/L. CFU were enumerated by serial dilutions avec centrifugation and resuspension of the pellets to avoid carry-over effect at 1, 3 and 24 hours. After 24 hours, the cultures were also plated on plates containing 4 times the MIC of ATM and 4 mg/mL AVI to look for resistant mutants. The detection limit was 1 colony per 100 µL.

### In vivo studies

#### Ethics and animal care

Animal experiments were performed in accordance with prevailing regulations regarding the care and use of laboratory animals from the European Commission. Experiments were approved by the local ethics committee (Departmental Direction of Veterinary Services, Paris France, agreement no. 22330-2019092415325730).

Swiss ICR-strain female mice weighing approximately 25 g were used. Animals were housed in regulations cages and given free access to food and water.

Overall, 253 mice were used for experiments.

#### Antibiotic pharmacokinetics

Antibiotics were injected subcutaneously to previously infected mice using the CFT073-pTOPO strain. ATM 100 mg/kg, AVI 100 mg/kg were administered separately or together in order to check for the absence of drug-drug interaction. Blood samples were obtained by intracardiac puncture from previously anesthetized mice (3 mice for each point), 15, 30, 60 and 120 minutes after injection of ATM or AVI, or the combination. Therapeutic regimens were chosen to achieve in serum a percentage of time during which the free drug concentrations exceeded the MIC (*%f*T _> MIC_) for ATM (considering MIC determined in standard conditions, according to the EUCAST recommendations, with a breakpoint of 1 mg/L for susceptibility), and ≥ 2.5 mg/L (%*f*T _CT > 2,5 mg/L_) for AVI, close to those obtained in humans with recommended dosages. The assays carried out by the laboratory yielded total fractions. We deduced the free fractions from protein binding in mice, assumed to be 43.5% for ATM and 10% for AVI^19^. Finally, we ran linear regression models to estimate *%f*T _> MIC_.

IPM was used for comparison because its activity is only slightly altered by the inoculum effect^33^. *In vivo*, a therapeutic regimen of 100 mg/kg/4h of IPM was chosen according to previous results of our team^28^.

#### Murine model of peritonitis

The murine model of peritonitis was used as previously described^34^. Briefly, overnight cultures of each strain were mixed with porcine mucin 10% (Sigma-Aldrich, Saint-Quentin-Fallavier, France). Mice were inoculated with an intraperitoneal injection of 250 µL of bacteria/mucin mix corresponding to a final inoculum of 10^6^ CFU (standard inoculum) of 10^8^ CFU (high inoculum). Two hours after inoculation, groups of 5 to 18 mice were treated with ATM, AVI, ATM plus AVI, or IPM for 24 hours. For each strain and inoculum, at least 3 mice per group were sacrificed 2 hours after inoculation to determine bacterial load (“start-of-treatment” (SOT) controls). In order to avoid animal suffering, animal well-being was evaluated every 4 hours then hourly if necessary according to protocol and mice were sacrificed if critical score was achieved^35^. Sacrifice was done by intraperitoneal injection of 400 mg/kg of sodium thiopental. Mice that survived during the whole duration of treatment were sacrificed 4 hours after the last antibiotic injection.

After death, *s*pleen was extracted and homogenized in 1 mL of sterile saline solution. Samples were plated onto agar containing kanamycin for quantitative culture. Results were expressed as log_10_ CFU/g for spleen. The detection limit was 1 log_10_ per gram.

### Statistical analysis

Results were expressed as median [minimum - maximum] for continuous variables. MGR and log_10_ CFU/g in spleen were compared for the different groups by the Mann-Whitney U-test or the Kruskal-Wallis test when appropriate. Death rate (mice sacrificed before end of protocol) in the different groups and area under the curve of the different pharmacokinetics assays were compared by the Fisher’s exact test. A p < 0.05 was considered significant.

## Results

### In vitro studies

#### Growth rate and fitness

MGRs were respectively 1.09 h^-1^ [0.99 – 1.22], 0.93 h^-1^ [0.89 – 1.15] and 0.82 h^-1^ [0.74 – 0.90] for *E. coli* CFT073-pTOPO, *E. coli* CFT073-NDM1 and *E. coli* CFT073-CTXM15-NDM1. MGR was significantly lower for the two strains carrying the NDM-1 plasmid compared to *E. coli* CFT073-pTOPO (p < 0.001). Also, MGR of *E. coli* CFT073-CTXM15-NDM1 was significantly lower than that of *E. coli* CFT073-NDM1 (p < 0.001).

#### MICs

MICs of ATM, AVI, ATM-AVI (4 mg/L) and IPM against each strain at standard and high inoculum are shown in Table 1.

**Table 1.** MICs of aztreonam, avibactam, aztreonam with a fixed concentration of avibactam 4 mg/L and imipenem determined by the broth microdilution method against *E. coli* strains at standard (10^5^ CFU/mL) and high (10^7^ CFU/mL) inoculum.

|  | MICs (mg/L) according to inoculum size (CFU/mL) |  |  |  |  |  |  |  |
| --- | --- | --- | --- | --- | --- | --- | --- | --- |
| *E. coli*<br>strain | ATM |  | AVI |  | ATM-AVI* |  | IPM |  |
| | $10^5$ | $10^7$ | $10^5$ | $10^7$ | $10^5$ | $10^7$ | $10^5$ | $10^7$ |
| **CFT073-**<br>**pTOPO** | 0.0625 | > 512 | 256 | > 512 | 0.0625 | 64 | 0.5 | 1 |
| **CFT073-**<br>**NDM1** | 0.0625 | > 512 | 256 | 512 | 0.031 | 64 | 256 | > 1024 |
| **CFT073-**<br>**CTXM15-**<br>**NDM1** | 256 | > 512 | 256 | > 512 | 0.031 | 64 | 64 | > 1024 |

\* fixed 4 mg/L concentration. ATM, aztreonam; AVI, avibactam; IPM, imipenem.

At standard (10^5^) inoculum, ATM was highly active against *E. coli* CFT073-pTOPO and *E. coli* CFT073-NDM1. CFT073-CTXM15-NDM1 was resistant to ATM but its activity was completely restored by the adjunction of AVI 4 mg/L. MICs of AVI against all tested strains were 256 mg/L, indicating the absence of significant intrinsic activity of AVI against these strains.

A major *in vitro* inoculum effect of ATM was evidenced, with MICs of ATM alone and of the ATM-AVI combination reaching > 512 and 64 mg/L respectively, at high (10^7^) inoculum, against all tested strains.

A major inoculum effect was also observed when MICs of ATM against *E. coli* CFT073-pTOPO were measured by the method of Steers *et al*^24^. Indeed, the MIC of ATM increased from 0.125 to 64 mg/L when increasing the inoculum from 10^5^ to 10^7^ CFU per spot, indicating that the observed inoculum effect was independent of the method used.

#### Impact of antibiotic exposure on bacterial morphology

Microscopic examination of bacteria recovered after 24h incubation with ATM, AVI and the combination at standard inoculum, is shown in Figure 1. Exposure of *E. coli* CFT073-pTOPO to ATM either alone or combined with AVI, and of *E. coli* CFT073-CTXM15-NDM1 to ATM/AVI resulted in bacterial filamentation. Exposure of *E. coli* CFT073-CTXM15-NDM1 to ATM alone did not result in bacterial filamentation. AVI alone had no effect on bacterial morphology. These results were the same with a standard or high inoculum (data for high inoculum not shown). Thus, ATM led to bacterial filamentation when active against the bacteria, i.e., in monotherapy or in combination with AVI against susceptible *E. coli* CFT073-pTOPO, and only in combination with AVI against *E. coli* CFT073-CTXM15-NDM1.

**Figure 1.**
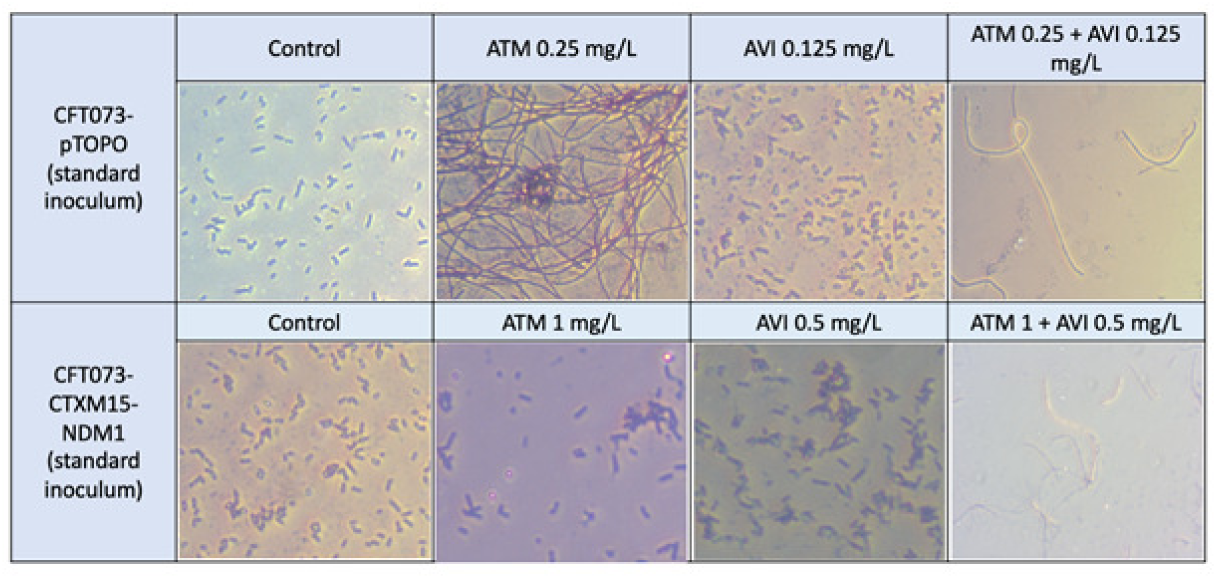
Gram stain morphology of *E. coli* CFT073-pTOPO and *E. coli* CFT073-CTXM15-NDM1 after 24h of incubation with ATM, AVI or ATM-AVI, or without antibiotic (control).

#### In vitro bactericidal activity of antibiotics and selection of mutants

Time kill curves of ATM 40 mg/L combined with AVI 4 mg/L against the three isolates at standard and high inoculum are shown in Figure 2. The concentration of AZT was chosen as it is equal to four times the free residual concentration in the infected mice treated with a single dose of 100 mg/kg AZT and thus seemed clinically relevant. The combination was bactericidal at 24h against the three isolates when a standard inoculum was used, but no bactericidal effect was observed at high inoculum. No mutants resistant to the combination were observed at 24h at low or high inoculum.

**Figure 2.**
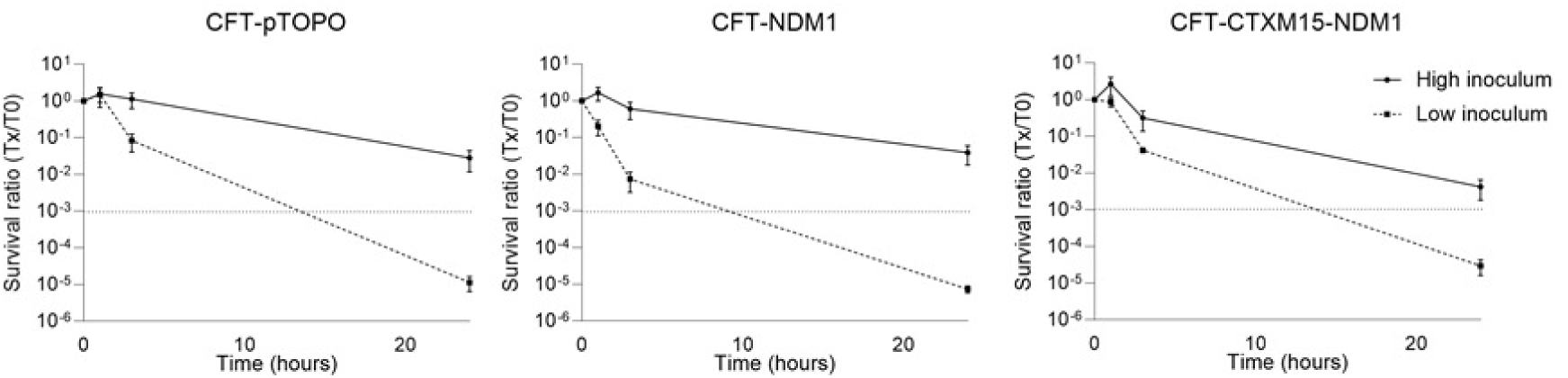
Time kill curves of Aztreonam 40 mg/L and avibactam 4 mg/L against *E. coli* CFT073-pTOPO, *E. coli* CFT073-NDM1 and *E. coli* CFT073-CTXM15-NDM1. Results are expressed as survival ratios (Tx/T0). High inoculum was between 5. 10^7^ and 2.10^8^ CFU/mL for *E. coli* CFT073-pTOPO, 1.10^8^ and 2.10^8^ CFU/mL for *E. coli* CFT073-NDM1 and 2.10^7^ and 7.10^7^ CFU/mL for *E. coli* CFT073-CTXM15-NDM1. The standard inoculum was one hundred times lower than the high inoculum.

### In vivo studies

#### Therapeutic regimen

Using a regimen of 100 mg/kg every 2h for ATM and AVI used in combination, the percentages of *f*T _> MIC_ (ATM) and *f*T _> 2.5 mg/L_ (AVI) were 100% and 95%, close to (ATM) or above (AVI) those obtained in humans at dosages recommended for severe infections (100% and 50%, respectively)^13^ (Table 2). There was no significant difference in dosages when drugs were administered separately or in combination (Figure 3). Indeed, the area under the curve (AUC) was not statistically different between ATM alone or in combination (86.1 and 63.3, respectively, p=0.31), nor between AVI alone or in combination (38.3 and 38.0, p=0.95).

**Figure 3.**
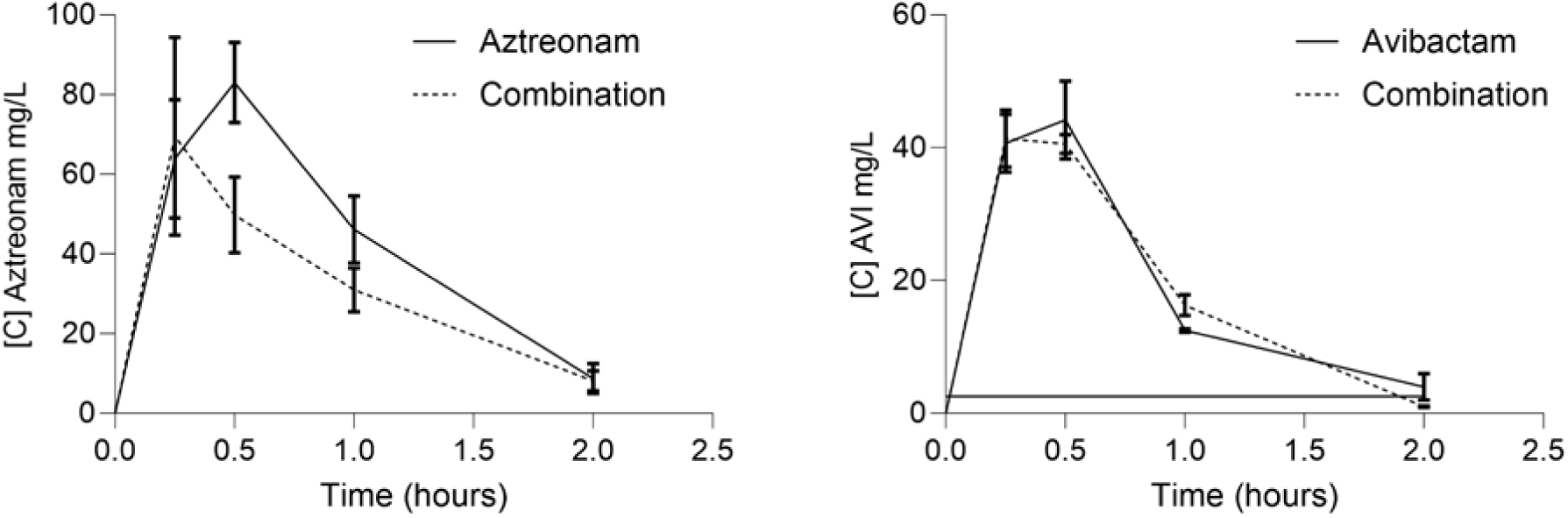
Pharmacokinetics of aztreonam and avibactam in infected mice. Female Swiss mice weighing 25 g were infected with an intraperitoneal injection of 10^6^ CFU of *E. coli* CFT073-pTOPO and treated two hours after infection with 100 mg/kg aztreonam or 100 mg/kg avibactam or a combination of both drugs. Mice were sacrificed 15, 30, 60 and 120 minutes after treatment and intracardiac puncture was performed. Graph represents free serum dose (HPLC) over time (n=3 to 6 mice per time point).

**Table 2.** Seric concentrations (total) and proportions of time during which free serum concentrations exceed the MIC (*(%f*T _> MIC_) for ATM and the critical threshold of 2.5 mg/L (*(%fT* _CT > 2,5 mg/L_) for AVI, according to therapeutic regimens used in humans^13^ and in mice infected with CFT073-pTOPO.

| Species | Antibiotic | ATM pharmacokinetic |  | AVI pharmacokinetic |  |
| --- | --- | --- | --- | --- | --- |
| | | Peak (mg/L) | $fT >_{MIC} (\%)*$ | Peak (mg/L) | $fT > 2.5 \text{ mg/L} (\%)$ |
| Mice | ATM 100 mg/kg | 158 ± 138 | 100% |  |  |
|  | AVI 100 mg/kg |  |  | 45 ± 8 | 95% |
| Humans | ATM 2 g | 105-242<sup>3,41</sup> | 100% |  |  |
|  | AVI 500 mg |  |  | 11,6-12,1<sup>25</sup> | 50% |

\**E. coli* CFT073-pTOPO and *E. coli* CFT073-CTXM15-NDM1 standard determination MICs
ATM, aztreonam; AVI, avibactam.

### Peritonitis

The efficacy of ATM and ATM/AVI in animals infected with *E. coli* CFT073-pTOPO and *E. coli* CFT073-CTXM15-NDM1, at standard and high inoculum is shown in Table 3. Full results are provided in the Supplementary Appendix (Table S1).

**Table 3.** Bacterial counts in spleen (log_10_ CFU/g), number of mice and survival rate in mice with peritonitis due *E. coli* CFT073-pTOPO and *E. coli* CFT073-CTXM15-NDM1 according to the inoculum size.

| | Median [range] $\log_{10}$ CFU/g of spleen<br>(number of surviving animals at sacrifice/number of total animals) | | | |
| --- | --- | --- | --- | --- |
| Regimen | Standard inoculum |  | High inoculum |  |
| *E. coli* strain | CFT073-pTOPO | CFT073-CTXM15-NDM1 | CFT073-pTOPO | CFT073-CTXM15-NDM1 |
| « Start-of-treatment » controls | 6.3 [5.9-6.4]<br>(3/3) | 3.9 [1.9-5.2]<br>(12/12) | 6.7 [4.3-8.0]<br>(6/6) | 6.8 [2.9-7.2]<br>(11/11) |
| ATM | 3.0 [2.5-3.2]*<br>(7/7) | 2.9 [2.1-3.5]<br>(14/14) | 6.3 [3.6-6.8]<br>(0/18)** | 5.9 [5.2-6.2]<br>(0/5)** |
| ATM + AVI | ND | 2.2 [1.7-2.8]*<br>(18/18) | ND | 5.3 [3.2-5.9]<br>(0/10)** |
| IPM | ND | ND | 4.3 [2.6-4.6]<br>(8/9)**** | ND |

ND, not done. ATM, aztreonam; AVI, avibactam; IPM, imipenem.
\* $p < 0.05$ versus « start of treatment » controls
\*\* $p < 0.01$ versus standard inoculum
\*\*\* $p < 0.01$ versus ATM at high inoculum

Using a standard inoculum, AVI alone had no activity against *E. coli* CFT073-CTXM15-NDM1 (death of 5/5 animals before end of treatment, median [range] log10 CFU/g of spleen after treatment, 4.4 [4.2-4.8], p = 0.29 versus SOT controls;). Based on this result and for ethical reasons, AVI alone was not tested at higher inoculum and/or against *E. coli* CFT073-pTOPO, since it was supposed to be inactive.

At standard inoculum, ATM alone was active in animals infected with *E. coli* CFT073-pTOPO both in terms of animal survival and bacterial decrease in spleens. In animals infected with *E. coli* CFT073-CTXM15-NDM1, ATM alone had no significant activity in term of spleen bacterial decrease by comparison to controls but led to bacterial survival; the combination of ATM and AVI displayed significant activity, with survival of 100% animals and a significant decrease in bacterial counts as compared to controls.

Increase in the inoculum led to a drastic decrease in antibiotic activity of ATM alone against *E. coli* CFT073-pTOPO, and of ATM-AVI against *E. coli* CFT073-CTXM15-NDM1. Indeed, using the same antibiotic regimens, all animals died and no significant reduction in spleen bacterial counts, as compared to controls, could be observed. At high inoculum, IPM was significantly active against *E. coli* CFT073-pTOPO.

Altogether, our results suggest a high impact of the inoculum increase on the activity of ATM against *E. coli* CFT073-pTOPO, and of the ATM-AVI combination against *E. coli* CFT073-CTXM15-NDM1 *in vivo*.

## Discussion

In the present study, we evidenced the *in vitro* activity of the ATM-AVI combination against CTX-M-15 plus NDM-1 producing *E. coli* in standard experimental conditions. ATM alone was effective against an *E. coli* CFT073-pTOPO and *E. coli* CFT073-NDM1 but inactive in the presence of the CTX-M-15 enzyme, as previously demonstrated^14^. AVI alone had no antibacterial activity against the 3 studied strains, however its presence was essential to restore the full activity of ATM against *E. coli* CFT073-CTXM15-NDM1^12^.

Exposition to a high bacterial inoculum led to a significant increase of the MICs of ATM rendering previously susceptible strains resistant to ATM according to EUCAST breakpoints^31^. This inoculum effect was independent of the method used for MIC determination. The explanations for the inoculum effect are not clearly defined, and many mechanisms may probably coexist^36^ . An explanation often put forward is the high amount of production of β-lactamases at high inoculum, that may overwhelm the amount of β-lactam (without and with a β-lactamase inhibitor); however, this mechanism is probably not the only one^37^ and does not explain the inoculum effect we observed with susceptible CFT073-pTOPO^34,38^. Another explanation may be the selection of pre-existing subpopulations resistant to ATM; of note, no mutants were retrieved at 24h when time kill curves were performed at high inoculum. The main explanation we propose for this loss of activity at high inoculum may be the filamentous growth of bacteria we observed, as others^33^, in the presence of ATM. Indeed, ATM binds exclusively penicillin-binding protein 3 (PBP3) which plays an important role in peptidoglycan synthesis during division process^39^. When PBP3 is inhibited, the segmentation of bacteria into bacilli is prevented, resulting in a filamentous growth of the strains. Accordingly, we observed a bacterial filamentation when ATM was incubated with bacteria, either alone or combined to AVI, provided that the bacteria was susceptible to the antibiotic used (ATM alone against *E. coli* CFT073-pTOPO, ATM-AVI combination against *E. coli* CFT073-CTXM15-NDM1). By contrast, no filamentation was observed when ATM was incubated with ATM-resistant *E. coli* CFT073-CTXM15-NDM1. Our results suggest that the inoculum effect may result from a change of the metabolic state of the bacteria, with a filamentation induced by ATM, which increases the bacterial biomass and density. At high inoculum, this increase in biomass is more noticeable than at low inoculum; it alters the susceptibility of these bacteria to ATM with or without AVI, which fails to reduce the number of viable organisms. Our results suggest that MIC determination, using the recommended EUCAST method with a standard inoculum of 5×10^5^ bacteria, could be inappropriate to predict ATM or ATM-AVI activity in high inoculum situations.

In the *in vivo* experiment, ATM permitted a significant decrease in bacterial count and the survival of all mice infected when animals were infected with a “standard” inoculum (inoculation of 10^6^ CFU) of *E. coli* CFT073-pTOPO. This result is in accordance with previously published data observed in the neutropenic murine thigh infection model^13^. As expected from *in vitro* results, ATM was inactive in term of decrease in bacterial counts against *E. coli* CFT073-CTXM15-NDM1. Interestingly, no animal death was observed over a short experiment of 24 hours, which may be explained by the loss of fitness of *E. coli* CFT073-CTXM15-NDM1, evidenced by an impaired *in vitro* growth rate. This significant resistance cost leading to a lower growth rate probably explains a lower infectivity, as shown by lower bacterial counts in SOT controls, and lower *in vivo* virulence, evidenced by animal survival.

It is important to assess the impact of the *in vitro* inoculum effect on the *in vivo* efficiency of antibiotic treatment, and the mouse model of peritonitis is a relevant model in this situation^34^. In fact, an *in vitro* inoculum effect doesn’t always lead to treatment failure *in vivo*. For example, using the same *in vivo* model, our team showed that despite a high inoculum effect *in vitro*, cefiderocol remained effective in a high inoculum peritonitis infection^40^. In the presence of a high inoculum (inoculation of 10^8^ CFU to animals), ATM against *E. coli* CFT073-pTOPO and ATM-AVI against *E. coli* CFT073-CTXM15-NDM1 were ineffective. Indeed, none of the mice in these groups survived until the end of protocol. It should be noted that therapeutic regimens used in mice led to concentrations close to those recommended in humans for ATM (*f*T _> MIC_ of 100%) but even higher than those recommended in humans for AVI (*f*T _> 2.5 mg/L_ of 95% in mice versus 50% in humans)^13^. In accordance with the *in vitro* results, the main hypothesis to explain this result could be a decreased ATM efficacy in the presence of a high inoculum. This hypothesis is backed up by the maintained activity of IPM against *E. coli* CFT073-pTOPO at high inoculum. Indeed, it was previously shown that IPM activity is poorly affected by the inoculum effect^33^. The impact of the inoculum on ATM has already been suggested by Soriano *et al*., reporting the need to drastically increase the dosage of antibiotics with a pronounced inoculum effect, such as ATM, to decrease mortality in a bacteriemic rats^6^. In the REJUVENATE study, a phase 2a open study evaluating the pharmacokinetics and safety of ATM plus AVI for the treatment of complicated intraabdominal infections, which presents the largest prospective cohort of patients treated with ATM-AVI, with 58.8% clinical cure rates at the test-of-cure visit, all patients had a surgical procedure for source control and peritoneal lavage responsible for a decreased inoculum^25^.

In conclusion, we show here the *in vivo* impact of the inoculum on the activity of ATM plus AVI against MBL-producing *Enterobacteriaceae*. ATM-AVI remains a promising option to treat infections caused by MBL-producing Enterobacteriaceae but treatment failures leading to deaths or recurrences were reported^21,23^. Our study suggests the implication of the inoculum effect in the onset of failures, although it cannot be demonstrated in clinical practice. Clinicians must be aware of the risk of failures when using ATM-AVI in high inoculum infections.

## Acknowledgements

None to declare.

## Funding

This study was supported by internal funding from “IAME”.

## Transparency declarations

None to declare.

## Notes

### Competing Interest Statement

The authors have declared no competing interest.

## References

1. Suay-García B, Pérez-Gracia MT. Present and Future of Carbapenem-resistant Enterobacteriaceae (CRE) Infections. Antibiotics (Basel) 2019;8(3):122.

2. Johnson DH, Cunha BA. Aztreonam. Med Clin North Am 1995;79(4):733–43.

3. Ramsey C, MacGowan AP. A review of the pharmacokinetics and pharmacodynamics of aztreonam. J Antimicrob Chemother 2016;71(10):2704–12.

4. Pulcini C, Bush K, Craig WA, et al. Forgotten antibiotics: an inventory in Europe, the United States, Canada, and Australia. Clin Infect Dis 2012;54(2):268–74.

5. Biedenbach DJ, Kazmierczak K, Bouchillon SK, Sahm DF, Bradford PA. In vitro activity of aztreonam-avibactam against a global collection of Gram-negative pathogens from 2012 and 2013. Antimicrob Agents Chemother 2015;59(7):4239–48.

6. Soriano F, Santamaría M, Ponte C, Castilla C, Fernández-Roblas R. In vivo significance of the inoculum effect of antibiotics on *Escherichia coli*. Eur J Clin Microbiol Infect Dis 1988;7(3):410–2.

7. Soriano F, Ponte C, Santamaría M, Jimenez-Arriero M. Relevance of the inoculum effect of antibiotics in the outcome of experimental infections caused by *Escherichia coli*. J Antimicrob Chemother 1990;25(4):621–7.

8. Goldstein EJ, Citron DM, Cherubin CE. Comparison of the inoculum effects of members of the family Enterobacteriaceae on cefoxitin and other cephalosporins, beta-lactamase inhibitor combinations, and the penicillin-derived components of these combinations. Antimicrob Agents Chemother 1991;35(3):560–6.

9. Nichols WW, Newell P, Critchley IA, Riccobene T, Das S. Avibactam Pharmacokinetic/Pharmacodynamic Targets. Antimicrob Agents Chemother 2018;62(6):e02446–17.

10. Ehmann DE, Jahic H, Ross PL, et al. Kinetics of avibactam inhibition against Class A, C, and D β-lactamases. J Biol Chem 2013;288(39):27960–71.

11. Ehmann DE, Jahić H, Ross PL, et al. Avibactam is a covalent, reversible, non-β-lactam β-lactamase inhibitor. Proc Natl Acad Sci U S A 2012;109(29):11663–8.

12. Berkhout J, Melchers MJ, van Mil AC, et al. Pharmacokinetics and penetration of ceftazidime and avibactam into epithelial lining fluid in thigh- and lung-infected mice. Antimicrob Agents Chemother 2015;59(4):2299–304.

13. Singh R, Kim A, Tanudra MA, et al. Pharmacokinetics/pharmacodynamics of a β-lactam and β-lactamase inhibitor combination: a novel approach for aztreonam/avibactam. J Antimicrob Chemother 2015;70(9):2618–26.

14. Livermore DM, Mushtaq S, Warner M, et al. Activities of NXL104 combinations with ceftazidime and aztreonam against carbapenemase-Producing Enterobacteriaceae. Antimicrob Agents Chemother 2011;55(1):390–4.

15. Huang Y-S, Chen P-Y, Chou P-C, Wang J-T. In Vitro Activities and Inoculum Effects of Cefiderocol and Aztreonam-Avibactam against Metallo-β-Lactamase-Producing Enterobacteriaceae. Microbiol Spectr 2023;11(3):e0056923.

16. Bae M, Kim T, Park JH, et al. In Vitro Activities of Ceftazidime–Avibactam and Aztreonam–Avibactam at Different Inoculum Sizes of Extended-Spectrum β-Lactam-Resistant Enterobacterales Blood Isolates. Antibiotics 2021;10(12):1492.

17. Danjean M, Hobson CA, Gits-Muselli M, et al. Evaluation of the inoculum effect of new antibiotics against carbapenem-resistant enterobacterales. Clin Microbiol Infect 2022;28(11):1503.e1–1503.e3.

18. Kim T, Lee SC, Bae M, et al. In Vitro Activities and Inoculum Effects of Ceftazidime-Avibactam and Aztreonam-Avibactam against Carbapenem-Resistant Enterobacterales Isolates from South Korea. Antibiotics (Basel) 2020;9(12):912.

19. Crandon JL, Nicolau DP. Human simulated studies of aztreonam and aztreonam-avibactam to evaluate activity against challenging gram-negative organisms, including metallo-β-lactamase producers. Antimicrob Agents Chemother 2013;57(7):3299–306.

20. Singh R, Kim A, Tanudra MA, et al. Pharmacokinetics/pharmacodynamics of a β-lactam and β-lactamase inhibitor combination: a novel approach for aztreonam/avibactam. Journal of Antimicrobial Chemotherapy 2015;70(9):2618–26.

21. Benchetrit L, Mathy V, Armand-Lefevre L, Bouadma L, Timsit J-F. Successful treatment of septic shock due to NDM-1-producing *Klebsiella pneumoniae* using ceftazidime/avibactam combined with aztreonam in solid organ transplant recipients: report of two cases. Int J Antimicrob Agents 2020;55(1):105842.

22. Hobson CA, Bonacorsi S, Fahd M, et al. Successful Treatment of Bacteremia Due to NDM-1-Producing *Morganella morganii* with Aztreonam and Ceftazidime-Avibactam Combination in a Pediatric Patient with Hematologic Malignancy. Antimicrob Agents Chemother 2019;63(2):e02463–18.

23. Shaw E, Rombauts A, Tubau F, et al. Clinical outcomes after combination treatment with ceftazidime/avibactam and aztreonam for NDM-1/OXA-48/CTX-M-15-producing *Klebsiella pneumoniae* infection. J Antimicrob Chemother 2018;73(4):1104–6.

24. Szymański M, Skiba MM, Piasecka M. Synergistic Effect of Ceftazidime-Avibactam with Aztreonam on Carbapenemase-Positive *Klebsiella pneumoniae* MBL+, NDM+ [Response to Letter]. Infection and Drug Resistance 2024;17:3159–60.

25. Cornely OA, Cisneros JM, Torre-Cisneros J, et al. Pharmacokinetics and safety of aztreonam/avibactam for the treatment of complicated intra-abdominal infections in hospitalized adults: results from the REJUVENATE study. J Antimicrob Chemother 2020;75(3):618–27.

26. Carmeli Y, Cisneros J-M, Paul M, et al. 2893 A. Efficacy and Safety of Aztreonam-Avibactam for the Treatment of Serious Infections Due to Gram-Negative Bacteria, Including Metallo-β-Lactamase-Producing Pathogens: Phase 3 REVISIT Study. Open Forum Infect Dis 2023;10(Suppl 2):ofad500.2476.

27. Tamma PD, Aitken SL, Bonomo RA, Mathers AJ, van Duin D, Clancy CJ. Infectious Diseases Society of America Guidance on the Treatment of Extended-Spectrum β-lactamase Producing Enterobacterales (ESBL-E), Carbapenem-Resistant Enterobacterales (CRE), and *Pseudomonas aeruginosa* with Difficult-to-Treat Resistance (DTR-P. aeruginosa). Clin Infect Dis 2021;72(7):e169–83.

28. Alexandre K, Chau F, Guérin F, et al. Activity of temocillin in a lethal murine model of infection of intra-abdominal origin due to KPC-producing *Escherichia coli*. J Antimicrob Chemother 2016;71(7):1899–904.

29. Welch RA, Burland V, Plunkett G, et al. Extensive mosaic structure revealed by the complete genome sequence of uropathogenic *Escherichia coli*. Proc Natl Acad Sci U S A 2002;99(26):17020–4.

30. Bleibtreu A, Gros P-A, Laouénan C, et al. Fitness, stress resistance, and extraintestinal virulence in *Escherichia coli*. Infect Immun 2013;81(8):2733–42.

31. eucast: Breakpoint tables and dosages v 12.0 (2022) published [Internet]. [cited 2024 Sep 16];Available from: https://www.eucast.org/eucast_news/news_singleview?tx_ttnews%5Btt_news%5D=464&cHash=ea8540c0fbdaa71b3bbcb3bf765239de

32. Steers E, Foltz EL, Graves BS. An inocula replicating apparatus for routine testing of bacterial susceptibility to antibiotics. Antibiot Chemother (Northfield) 1959;9(5):307–11.

33. Eng RH, Cherubin C, Smith SM, Buccini F. Inoculum effect of beta-lactam antibiotics on Enterobacteriaceae. Antimicrob Agents Chemother 1985;28(5):601–6.

34. Docobo-Pérez F, López-Cerero L, López-Rojas R, et al. Inoculum effect on the efficacies of amoxicillin-clavulanate, piperacillin-tazobactam, and imipenem against extended-spectrum β-lactamase (ESBL)-producing and non-ESBL-producing *Escherichia coli* in an experimental murine sepsis model. Antimicrob Agents Chemother 2013;57(5):2109–13.

35. Shrum B, Anantha RV, Xu SX, et al. A robust scoring system to evaluate sepsis severity in an animal model. BMC Res Notes 2014;7:233.

36. Lenhard JR, Bulman ZP. Inoculum effect of β-lactam antibiotics. J Antimicrob Chemother 2019;74(10):2825–43.

37. López-Cerero L, Picón E, Morillo C, et al. Comparative assessment of inoculum effects on the antimicrobial activity of amoxycillin-clavulanate and piperacillin-tazobactam with extended-spectrum beta-lactamase-producing and extended-spectrum beta-lactamase-non-producing *Escherichia coli* isolates. Clin Microbiol Infect 2010;16(2):132–6.

38. Bulger RR, Washington JA. Effect of inoculum size and beta-lactamase production on in vitro activity of new cephalosporins against *Haemophilus* species. Antimicrob Agents Chemother 1980;17(3):393–6.

39. Nguyen-Distèche M, Fraipont C, Buddelmeijer N, Nanninga N. The structure and function of *Escherichia coli* penicillin-binding protein 3. Cell Mol Life Sci 1998;54(4):309– 16.

40. Godron AS, Amoura A, Pistien C, Magréault S, Jousset A, Dion S, Jullien V, Lefort A, Fantin B, El Meouche I, de Lastours V. Cefiderocol and inoculum effect : discordant results between *in vitro* and murine model of peritonitis2024. Abstr 24th European Congress of clinical Microbiology and Infectious Diseases, abstr P3528.

41. Swabb EA, Leitz MA, Pilkiewicz FG, Sugerman AA. Pharmacokinetics of the monobactam SQ 26,776 after single intravenous doses in healthy subjects. J Antimicrob Chemother 1981;8 Suppl E:131–40.

