## Supplementary Table 1 for "Impact of the inoculum size on the *in vivo* activity of the aztreonam-avibactam combination in a murine model of peritonitis due to *Escherichia coli* expressing CTX-M-15 and NDM-1"

**Supplementary Appendix**

Table S1. Individual results of mice infected with *E. coli* CFT073-pTOPO and *E. coli* CFT073-CTXM15-NDM1 and treated with .

| **Mouse #** | **Strain** | **Inoculum** | **Bacterial count** | **Regimen** | **Bacterial count (log_10_ CFU/g of spleen)** | **Survival at sacrifice** | **Comment** |
| --- | --- | --- | --- | --- | --- | --- | --- |
| 35 | CFT073-pTOPO | Standard | 8,00E+06 | Start-of-treatment (SOT) control | 5,9 | Yes |  |
| 36 | CFT073-pTOPO | Standard | 8,00E+06 | SOT control | 6,4 | Yes |  |
| 37 | CFT073-pTOPO | Standard | 8,00E+06 | SOT control | 6,3 | Yes |  |
| 41 | CFT073-pTOPO | Standard | 8,00E+06 | Aztreonam (ATM) 100 mg/kg q2h | 2,1 | Yes | Not infected - excluded |
| 42 | CFT073-pTOPO | Standard | 8,00E+06 | ATM 100 mg/kg q2h | 2,8 | Yes |  |
| 43 | CFT073-pTOPO | Standard | 8,00E+06 | ATM 100 mg/kg q2h | 3,1 | Yes |  |
| 44 | CFT073-pTOPO | Standard | 8,00E+06 | ATM 100 mg/kg q2h | 3,0 | Yes |  |
| 45 | CFT073-pTOPO | Standard | 8,00E+06 | ATM 100 mg/kg q2h | 2,5 | Yes |  |
| 46 | CFT073-pTOPO | Standard | 8,00E+06 | ATM 100 mg/kg q2h | 2,7 | Yes |  |
| 47 | CFT073-pTOPO | Standard | 8,00E+06 | ATM 100 mg/kg q2h | 3,2 | Yes |  |
| 48 | CFT073-pTOPO | Standard | 8,00E+06 | ATM 100 mg/kg q2h | 3,2 | Yes |  |
| 1 | CFT073-pTOPO | High | 6,00E+08 | SOT control | 6,5 | Yes |  |
| 2 | CFT073-pTOPO | High | 6,00E+08 | SOT control | 7,0 | Yes |  |
| 3 | CFT073-pTOPO | High | 6,00E+08 | SOT control | 5,6 | Yes |  |
| 38 | CFT073-pTOPO | High | 5,40E+08 | SOT control | 7,2 | Yes |  |
| 39 | CFT073-pTOPO | High | 5,40E+08 | SOT control | 4,3 | Yes |  |
| 40 | CFT073-pTOPO | High | 5,40E+08 | SOT controls | 8,0 | Yes |  |
| 10 | CFT073-pTOPO | High | 6,00E+08 | ATM 100 mg/kg q2h | 5,8 | No | Sacrifice before end of protocol |
| 11 | CFT073-pTOPO | High | 6,00E+08 | ATM 100 mg/kg q2h | 6,6 | No | Sacrifice before end of protocol |
| 12 | CFT073-pTOPO | High | 6,00E+08 | ATM 100 mg/kg q2h | 6,2 | No | Sacrifice before end of protocol |
| 13 | CFT073-pTOPO | High | 6,00E+08 | ATM 100 mg/kg q2h | 6,1 | No | Sacrifice before end of protocol |
| 14 | CFT073-pTOPO | High | 6,00E+08 | ATM 100 mg/kg q2h | 5,7 | No | Sacrifice before end of protocol |
| 15 | CFT073-pTOPO | High | 6,00E+08 | ATM 100 mg/kg q2h | 6,2 | No | Sacrifice before end of protocol |
| 16 | CFT073-pTOPO | High | 6,00E+08 | ATM 100 mg/kg q2h | 3,6 | No | Sacrifice before end of protocol |
| 17 | CFT073-pTOPO | High | 6,00E+08 | ATM 100 mg/kg q2h | 5,1 | No | Sacrifice before end of protocol |
| 18 | CFT073-pTOPO | High | 6,00E+08 | ATM 100 mg/kg q2h | 6,8 | No | Sacrifice before end of protocol |
| 19 | CFT073-pTOPO | High | 6,00E+08 | ATM 100 mg/kg q2h | 6,0 | No | Sacrifice before end of protocol |
| 49 | CFT073-pTOPO | High | 5,40E+08 | ATM 100 mg/kg q2h | 6,4 | No | Sacrifice before end of protocol |
| 50 | CFT073-pTOPO | High | 5,40E+08 | ATM 100 mg/kg q2h | 6,1 | No | Sacrifice before end of protocol |
| 51 | CFT073-pTOPO | High | 5,40E+08 | ATM 100 mg/kg q2h | 6,6 | No | Sacrifice before end of protocol |
| 52 | CFT073-pTOPO | High | 5,40E+08 | ATM 100 mg/kg q2h | 6,5 | No | Sacrifice before end of protocol |
| 53 | CFT073-pTOPO | High | 5,40E+08 | ATM 100 mg/kg q2h | 6,5 | No | Sacrifice before end of protocol |
| 54 | CFT073-pTOPO | High | 5,40E+08 | ATM 100 mg/kg q2h | 6,5 | No | Sacrifice before end of protocol |
| 55 | CFT073-pTOPO | High | 5,40E+08 | ATM 100 mg/kg q2h | 6,3 | No | Sacrifice before end of protocol |
| 56 | CFT073-pTOPO | High | 5,40E+08 | ATM 100 mg/kg q2h | 6,3 | No | Sacrifice before end of protocol |
| 57 | CFT073-pTOPO | High | 5,40E+08 | Imipenem (IPM) 100 mg/kg q4h | 4,2 | Yes |  |
| 58 | CFT073-pTOPO | High | 5,40E+08 | IPM 100 mg/kg q4h | 4,5 | Yes |  |
| 59 | CFT073-pTOPO | High | 5,40E+08 | IPM 100 mg/kg q4h | 4,3 | Yes |  |
| 60 | CFT073-pTOPO | High | 5,40E+08 | IPM 100 mg/kg q4h | 4,0 | Yes |  |
| 61 | CFT073-pTOPO | High | 5,40E+08 | IPM 100 mg/kg q4h | 4,3 | Yes |  |
| 62 | CFT073-pTOPO | High | 5,40E+08 | IPM 100 mg/kg q4h | 4,4 | Yes |  |
| 63 | CFT073-pTOPO | High | 5,40E+08 | IPM 100 mg/kg q4h | 2,6 | Yes |  |
| 64 | CFT073-pTOPO | High | 5,40E+08 | IPM 100 mg/kg q4h | 4,6 | Yes |  |
| 65 | CFT073-pTOPO | High | 5,40E+08 | IPM 100 mg/kg q4h | 3,9 | No | Sacrifice before end of protocol |
| 66 | CFT073-CTXM15-NDM1 | Standard | 4,80E+06 | SOT control | 4,8 | Yes |  |
| 67 | CFT073-CTXM15-NDM1 | Standard | 4,80E+06 | SOT control | 2,5 | Yes |  |
| 68 | CFT073-CTXM15-NDM1 | Standard | 4,80E+06 | SOT control | 2,7 | Yes |  |
| 69 | CFT073-CTXM15-NDM1 | Standard | 4,80E+06 | SOT control | 5,0 | Yes |  |
| 70 | CFT073-CTXM15-NDM1 | Standard | 4,80E+06 | SOT control | 4,7 | Yes |  |
| 104 | CFT073-CTXM15-NDM1 | Standard | 1,60E+06 | SOT control | 2,3 | Yes |  |
| 105 | CFT073-CTXM15-NDM1 | Standard | 1,60E+06 | SOT control | 3,9 | Yes |  |
| 106 | CFT073-CTXM15-NDM1 | Standard | 1,60E+06 | SOT control | 1,9 | Yes |  |
| 107 | CFT073-CTXM15-NDM1 | Standard | 1,60E+06 | SOT control | 3,9 | Yes |  |
| 108 | CFT073-CTXM15-NDM1 | Standard | 1,60E+06 | SOT control | 3,9 | Yes |  |
| 109 | CFT073-CTXM15-NDM1 | Standard | 1,60E+06 | SOT control | 4,0 | Yes |  |
| 110 | CFT073-CTXM15-NDM1 | Standard | 1,60E+06 | SOT control | 1,8 | Yes | Not infected - Excluded |
| 111 | CFT073-CTXM15-NDM1 | Standard | 1,60E+06 | SOT control | 5,2 | Yes |  |
| 74 | CFT073-CTXM15-NDM1 | Standard | 4,80E+06 | ATM 100 mg/kg q2h | 2,2 | Yes |  |
| 75 | CFT073-CTXM15-NDM1 | Standard | 4,80E+06 | ATM 100 mg/kg q2h | 3,1 | Yes |  |
| 76 | CFT073-CTXM15-NDM1 | Standard | 4,80E+06 | ATM 100 mg/kg q2h | 2,9 | Yes |  |
| 77 | CFT073-CTXM15-NDM1 | Standard | 4,80E+06 | ATM 100 mg/kg q2h | 3,2 | Yes |  |
| 112 | CFT073-CTXM15-NDM1 | Standard | 1,60E+06 | ATM 100 mg/kg q2h | 3,5 | Yes |  |
| 113 | CFT073-CTXM15-NDM1 | Standard | 1,60E+06 | ATM 100 mg/kg q2h | 2,8 | Yes |  |
| 114 | CFT073-CTXM15-NDM1 | Standard | 1,60E+06 | ATM 100 mg/kg q2h | 2,5 | Yes |  |
| 115 | CFT073-CTXM15-NDM1 | Standard | 1,60E+06 | ATM 100 mg/kg q2h | 2,9 | Yes |  |
| 116 | CFT073-CTXM15-NDM1 | Standard | 1,60E+06 | ATM 100 mg/kg q2h | 2,7 | Yes |  |
| 117 | CFT073-CTXM15-NDM1 | Standard | 1,60E+06 | ATM 100 mg/kg q2h | 2,1 | Yes |  |
| 118 | CFT073-CTXM15-NDM1 | Standard | 1,60E+06 | ATM 100 mg/kg q2h | 3,4 | Yes |  |
| 119 | CFT073-CTXM15-NDM1 | Standard | 1,60E+06 | ATM 100 mg/kg q2h | 2,4 | Yes |  |
| 120 | CFT073-CTXM15-NDM1 | Standard | 1,60E+06 | ATM 100 mg/kg q2h | 2,5 | Yes |  |
| 121 | CFT073-CTXM15-NDM1 | Standard | 1,60E+06 | ATM 100 mg/kg q2h | 2,9 | Yes |  |
| 84 | CFT073-CTXM15-NDM1 | Standard | 4,80E+06 | ATM 100 mg/kg q2h + Avibactam (AVI) 100 mg/kg q2h | 2,3 | Yes |  |
| 85 | CFT073-CTXM15-NDM1 | Standard | 4,80E+06 | ATM 100 mg/kg q2h + AVI 100 mg/kg q2h | 2,2 | Yes |  |
| 86 | CFT073-CTXM15-NDM1 | Standard | 4,80E+06 | ATM 100 mg/kg q2h + AVI 100 mg/kg q2h | 0,9 | Yes | Not infected - Excluded |
| 87 | CFT073-CTXM15-NDM1 | Standard | 4,80E+06 | ATM 100 mg/kg q2h + AVI 100 mg/kg q2h | 2,1 | Yes |  |
| 88 | CFT073-CTXM15-NDM1 | Standard | 4,80E+06 | ATM 100 mg/kg q2h + AVI 100 mg/kg q2h | 2,1 | Yes |  |
| 89 | CFT073-CTXM15-NDM1 | Standard | 4,80E+06 | ATM 100 mg/kg q2h + AVI 100 mg/kg q2h | 2,8 | Yes |  |
| 90 | CFT073-CTXM15-NDM1 | Standard | 4,80E+06 | ATM 100 mg/kg q2h + AVI 100 mg/kg q2h | 2,1 | Yes |  |
| 91 | CFT073-CTXM15-NDM1 | Standard | 4,80E+06 | ATM 100 mg/kg q2h + AVI 100 mg/kg q2h | 2,5 | Yes |  |
| 92 | CFT073-CTXM15-NDM1 | Standard | 4,80E+06 | ATM 100 mg/kg q2h + AVI 100 mg/kg q2h | 2,7 | Yes |  |
| 93 | CFT073-CTXM15-NDM1 | Standard | 4,80E+06 | ATM 100 mg/kg q2h + AVI 100 mg/kg q2h | 0,9 | Yes | Not infected - Excluded |
| 122 | CFT073-CTXM15-NDM1 | Standard | 1,60E+06 | ATM 100 mg/kg q2h + AVI 100 mg/kg q2h | 2,1 | Yes |  |
| 123 | CFT073-CTXM15-NDM1 | Standard | 1,60E+06 | ATM 100 mg/kg q2h + AVI 100 mg/kg q2h | 2,4 | Yes |  |
| 124 | CFT073-CTXM15-NDM1 | Standard | 1,60E+06 | ATM 100 mg/kg q2h + AVI 100 mg/kg q2h | 1,8 | Yes |  |
| 125 | CFT073-CTXM15-NDM1 | Standard | 1,60E+06 | ATM 100 mg/kg q2h + AVI 100 mg/kg q2h | 2,3 | Yes |  |
| 126 | CFT073-CTXM15-NDM1 | Standard | 1,60E+06 | ATM 100 mg/kg q2h + AVI 100 mg/kg q2h | 1,7 | Yes |  |
| 127 | CFT073-CTXM15-NDM1 | Standard | 1,60E+06 | ATM 100 mg/kg q2h + AVI 100 mg/kg q2h | 2,3 | Yes |  |
| 128 | CFT073-CTXM15-NDM1 | Standard | 1,60E+06 | ATM 100 mg/kg q2h + AVI 100 mg/kg q2h | 2,6 | Yes |  |
| 129 | CFT073-CTXM15-NDM1 | Standard | 1,60E+06 | ATM 100 mg/kg q2h + AVI 100 mg/kg q2h | 1,9 | Yes |  |
| 130 | CFT073-CTXM15-NDM1 | Standard | 1,60E+06 | ATM 100 mg/kg q2h + AVI 100 mg/kg q2h | 1,7 | Yes |  |
| 131 | CFT073-CTXM15-NDM1 | Standard | 1,60E+06 | ATM 100 mg/kg q2h + AVI 100 mg/kg q2h | 1,9 | Yes |  |
| 7 | CFT073-CTXM15-NDM1 | High | 2,00E+08 | SOT control | 6,8 | Yes |  |
| 8 | CFT073-CTXM15-NDM1 | High | 2,00E+08 | SOT control | 6,8 | Yes |  |
| 9 | CFT073-CTXM15-NDM1 | High | 2,00E+08 | SOT control | 6,7 | Yes |  |
| 71 | CFT073-CTXM15-NDM1 | High | 2,40E+08 | SOT control | 3,1 | Yes |  |
| 72 | CFT073-CTXM15-NDM1 | High | 2,40E+08 | SOT control | 3,1 | Yes |  |
| 73 | CFT073-CTXM15-NDM1 | High | 2,40E+08 | SOT control | 6,8 | Yes |  |
| 132 | CFT073-CTXM15-NDM1 | High | 2,80E+08 | SOT control | 2,9 | Yes |  |
| 133 | CFT073-CTXM15-NDM1 | High | 2,80E+08 | SOT control | 6,8 | Yes |  |
| 134 | CFT073-CTXM15-NDM1 | High | 2,80E+08 | SOT control | 7,2 | Yes |  |
| 135 | CFT073-CTXM15-NDM1 | High | 2,80E+08 | SOT control | 7,0 | Yes |  |
| 136 | CFT073-CTXM15-NDM1 | High | 2,80E+08 | SOT control | 7,1 | Yes |  |
| 30 | CFT073-CTXM15-NDM1 | High | 2,00E+08 | ATM 100 mg/kg q2h | 6,2 | No | Sacrifice before end of protocol |
| 31 | CFT073-CTXM15-NDM1 | High | 2,00E+08 | ATM 100 mg/kg q2h | 6,1 | No | Sacrifice before end of protocol |
| 32 | CFT073-CTXM15-NDM1 | High | 2,00E+08 | ATM 100 mg/kg q2h | 5,9 | No | Sacrifice before end of protocol |
| 33 | CFT073-CTXM15-NDM1 | High | 2,00E+08 | ATM 100 mg/kg q2h | 5,2 | No | Sacrifice before end of protocol |
| 34 | CFT073-CTXM15-NDM1 | High | 2,00E+08 | ATM 100 mg/kg q2h | 5,3 | No | Sacrifice before end of protocol |
| 94 | CFT073-CTXM15-NDM1 | High | 2,40E+08 | ATM 100 mg/kg q2h + AVI 100 mg/kg q2h | 5,4 | No | Sacrifice before end of protocol |
| 95 | CFT073-CTXM15-NDM1 | High | 2,40E+08 | ATM 100 mg/kg q2h + AVI 100 mg/kg q2h | 5,6 | No | Sacrifice before end of protocol |
| 96 | CFT073-CTXM15-NDM1 | High | 2,40E+08 | ATM 100 mg/kg q2h + AVI 100 mg/kg q2h | 4,8 | No | Sacrifice before end of protocol |
| 97 | CFT073-CTXM15-NDM1 | High | 2,40E+08 | ATM 100 mg/kg q2h + AVI 100 mg/kg q2h | 5,3 | No | Sacrifice before end of protocol |
| 98 | CFT073-CTXM15-NDM1 | High | 2,40E+08 | ATM 100 mg/kg q2h + AVI 100 mg/kg q2h | 5,4 | No | Sacrifice before end of protocol |
| 99 | CFT073-CTXM15-NDM1 | High | 2,40E+08 | ATM 100 mg/kg q2h + AVI 100 mg/kg q2h | 5,4 | No | Sacrifice before end of protocol |
| 100 | CFT073-CTXM15-NDM1 | High | 2,40E+08 | ATM 100 mg/kg q2h + AVI 100 mg/kg q2h | 5,1 | No | Sacrifice before end of protocol |
| 101 | CFT073-CTXM15-NDM1 | High | 2,40E+08 | ATM 100 mg/kg q2h + AVI 100 mg/kg q2h | 5,9 | No | Sacrifice before end of protocol |
| 102 | CFT073-CTXM15-NDM1 | High | 2,40E+08 | ATM 100 mg/kg q2h + AVI 100 mg/kg q2h | 3,2 | No | Sacrifice before end of protocol |
| 103 | CFT073-CTXM15-NDM1 | High | 2,40E+08 | ATM 100 mg/kg q2h + AVI 100 mg/kg q2h | 3,2 | No | Sacrifice before end of protocol |

ATM, aztreonam; AVI, avibactam; SOT, start-of-treatment
